# Genome-scale metabolic model of *Staphylococcus epidermidis* ATCC 12228 matches *in vitro* conditions

**DOI:** 10.1101/2023.12.19.572329

**Authors:** Nantia Leonidou, Alina Renz, Benjamin Winnerling, Anastasiia Grekova, Fabian Grein, Andreas Dräger

## Abstract

*Staphylococcus epidermidis*, a commensal bacterium inhabiting collagen-rich areas, like human skin, has gained significance due to its probiotic potential in the nasal microbiome and as a leading cause of nosocomial infections. While infrequently leading to severe illnesses, *S. epidermidis* exerts a significant influence, particularly in its close association with implant-related infections and its role as a classic opportunistic biofilm former. Understanding its opportunistic nature is crucial for developing novel therapeutic strategies, addressing both its beneficial and pathogenic aspects, and alleviating the burdens it imposes on patients and healthcare systems. Here, we employ genome-scale metabolic modeling as a powerful tool to elucidate the lifestyle and capabilities of *S. epidermidis*. We created a comprehensive computational resource for understanding the organism’s growth conditions within diverse habitats by reconstructing and analyzing a manually curated and experimentally validated metabolic model. The final network, *i*Sep23, incorporates 1,415 reactions, 1,051 metabolites, and 705 genes, adhering to established community standards and modeling guidelines. Benchmarking with the `MEMOTE` test suite yields a high score, highlighting the model’s high semantic quality. Following the FAIR data principles, *i*Sep23 becomes a valuable and publicly accessible asset for subsequent studies. Growth simulations and carbon source utilization predictions align with experimental results, showcasing the model’s predictive power. This metabolic model advances our understanding of *S. epidermidis* as a commensal and potential probiotic and enhances insights into its opportunistic pathogenicity against other microorganisms.

**Author summary:** *Staphylococcus epidermidis*, a bacterium commonly found on human skin, has shown probiotic effects in the nasal microbiome and is a notable causative agent of hospital-acquired infections. While typically causing non-life-threatening diseases, the economic ramifications of *S. epidermidis* infections are substantial, with annual costs reaching billions of dollars in the United States. To unravel its opportunistic nature, we utilized genome-scale metabolic modeling, creating a detailed mathematical network that elucidates *S. epidermidis*’s lifestyle and capabilities. This model, encompassing over a thousand reactions, metabolites, and genes, adheres rigorously to established standards and guidelines, evident in its commendable benchmarking scores. Adhering to the FAIR data principles (Findable, Accessible, Interoperable, and Reusable), the model stands as a valuable resource for subsequent investigations. Growth simulations and predictions align closely with experimental results, showcasing the model’s predictive accuracy. This metabolic model not only enhances our understanding of *S. epidermidis* as a skin commensal and potential probiotic but also sheds light on its opportunistic pathogenicity, particularly in competition with other microorganisms.

## Introduction

A prevalent constituent of the human skin flora is the coagulase-negative commensal *Staphylococcus epidermidis*^1, 2^. This Gram-positive coccus predominantly inhabits the skin and mucosal membranes in areas such as the axillae, head, legs, arms, and nares. *S. epidermidis* plays a crucial role in maintaining a balanced microbiome within the human nasal cavity, where harmful pathogens like *Staphylococcus aureus* commonly establish colonization. There is ongoing discourse regarding whether *S. epidermidis*, through competition in nutritionally scarce environments like the human nose, may exhibit probiotic effects against formidable pathogens such as *S. aureus*^2, 3^. Nevertheless, *S. epidermidis* is recognized as a significant causative agent of nosocomial infections under specific conditions^4^. Notably, *S. epidermidis* stands out as the primary source of infections associated with indwelling medical devices, including intravascular catheters and implants such as prosthetic joints^1, 5, 6^. The high occurrence of these nosocomial infections is attributed to *S. epidermidis*’s ubiquitous presence on the human skin, increasing the likelihood of contamination during the insertion of medical devices^7^. Upon infection, *S. epidermidis* strains are capable of forming biofilms that shield them from antibiotics and host defense mechanisms, rendering *S. epidermidis* infections resistant and challenging to eliminate^1, 7^. Often, removing the foreign material becomes necessary to combat the infection effectively. While *S. epidermidis* infections seldom lead to life-threatening conditions, their impact on patients and the public health system is substantial. In the United States alone, the annual economic burden of *S. epidermidis* vascular catheter-related bloodstream infections is estimated to be around $2 billion^1, 6^. Besides biofilm formation, also other specific molecular determinants contribute to the pathogenicity of this particular pathogen, enabling immune evasion. Therefore, there is an urgent need for a more comprehensive understanding of *S. epidermidis* and its opportunistic characteristics to identify novel therapeutic strategies^1, 7^.

One way to better understand an organism’s lifestyle and capabilities is the reconstruction and analysis of genome-scale metabolic models (GEMs). These models rely on the annotated genome sequence of the organism in question. Specifically, genes encoding proteins with metabolic significance are allocated to their respective reactions through gene-protein-reaction associations (GPRs). Within the resulting network, biochemical reactions establish connections between metabolites, with enzymatic activities guided by genes associated with these reactions. Such models enable the comprehension of an organism’s metabolism at a systemic level. Díaz Calvo et al. reconstructed the metabolic network of RP62A, a slime-producing and methicillin-resistant biofilm isolate^8^. However, the resulted model is available only upon request. Figure 1 summarizes the computational and experimental approach of this article. This work introduces *i*Sep23, the first manually curated and experimentally validated GEM of *S. epidermidis* ATCC 12228. The model comprises 1,415 reactions, 1,051 metabolites, and 705 genes and is freely available from BioModels Database^9^ with the accession identifier `MODEL2012220002`. Moreover, it aligns with current community standards^10, 11, 12^ and modeling guidelines^13, 14^. Semantic benchmarking was conducted utilizing the `MEMOTE` genome-scale metabolic model test suite^15^. Consequently, *i*Sep23 upholds the Findable, Accessible, Interoperable, and Reusable (FAIR) data principles^16^, rendering it a valuable resource for subsequent research^17, 18^. To assess the predictive capacity of the model, growth simulations in various media were compared against laboratory experiments. The model’s predictions regarding the utilization of diverse carbon sources were cross-referenced with experimental findings. Altogether, our model establishes a foundation for improved comprehension of the organism’s phenotypes and behavior under different nutritional conditions.

**Figure 1.**
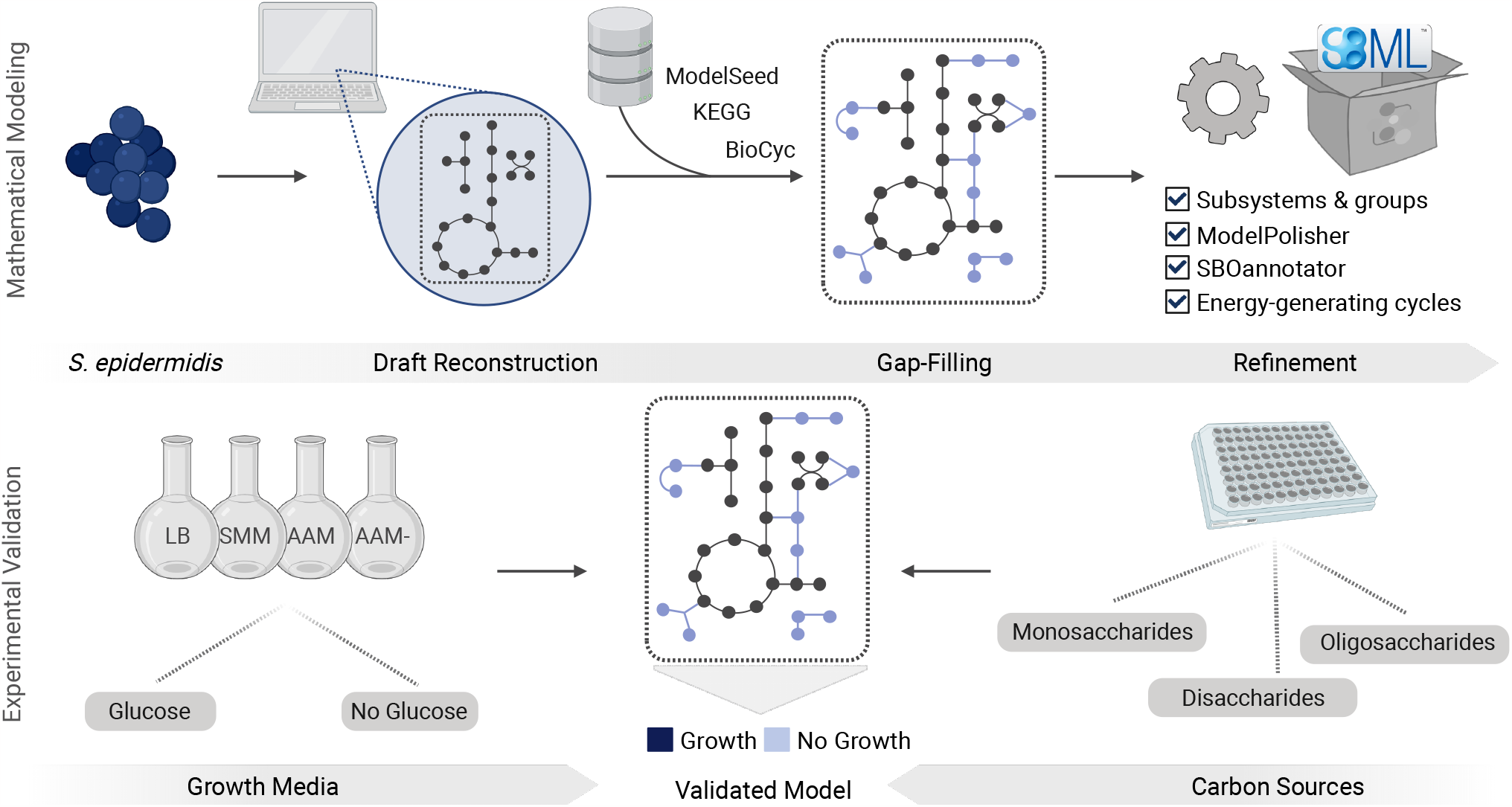
A new metabolic network for *S. epidermidis* ATCC 12228, called *i*Sep23. The computational metabolic network was created and validated using a two-phase approach. The initial phase encompassed the mathematical representation of the metabolism through the deployment of genome-scale models. Subsequently, the second phase involved the functional validation, rooted in experimental data.

## Results

### Properties of the constructed GEM

The initial CarveMe draft comprised 1,295 reactions, 933 metabolites, and 722 genes, yielding a Metabolic Model Testing (MEMOTE) ^15^ score of 36 %. Subsequent manual refinement involved the addition of 120 reactions, 118 metabolites, and 63 genes, as illustrated in Figure 2, resulting in an overall MEMOTE score of 88 %. The 63 mass- and charge-imbalanced reactions were reduced to one mass-imbalanced and nine charge-imbalanced reactions, resulting in a MEMOTE mass balance score of 99.7 % and a charge balance score of 99.3 %. Based on literature evidence, we corrected the directionality of 34 enzymatic reactions in the model to ensure proper constraints during model simulations. Moreover, the final metabolic network does not include infeasible energy generating cycle (EGC) that could inflate the simulation results (see Materials and Methods). We annotated the model instances with cross-references to various databases and additional information to increase the model’s interoperability and re-usability. The reaction annotations are divided into three different biological qualifier types:

**Figure 2.**
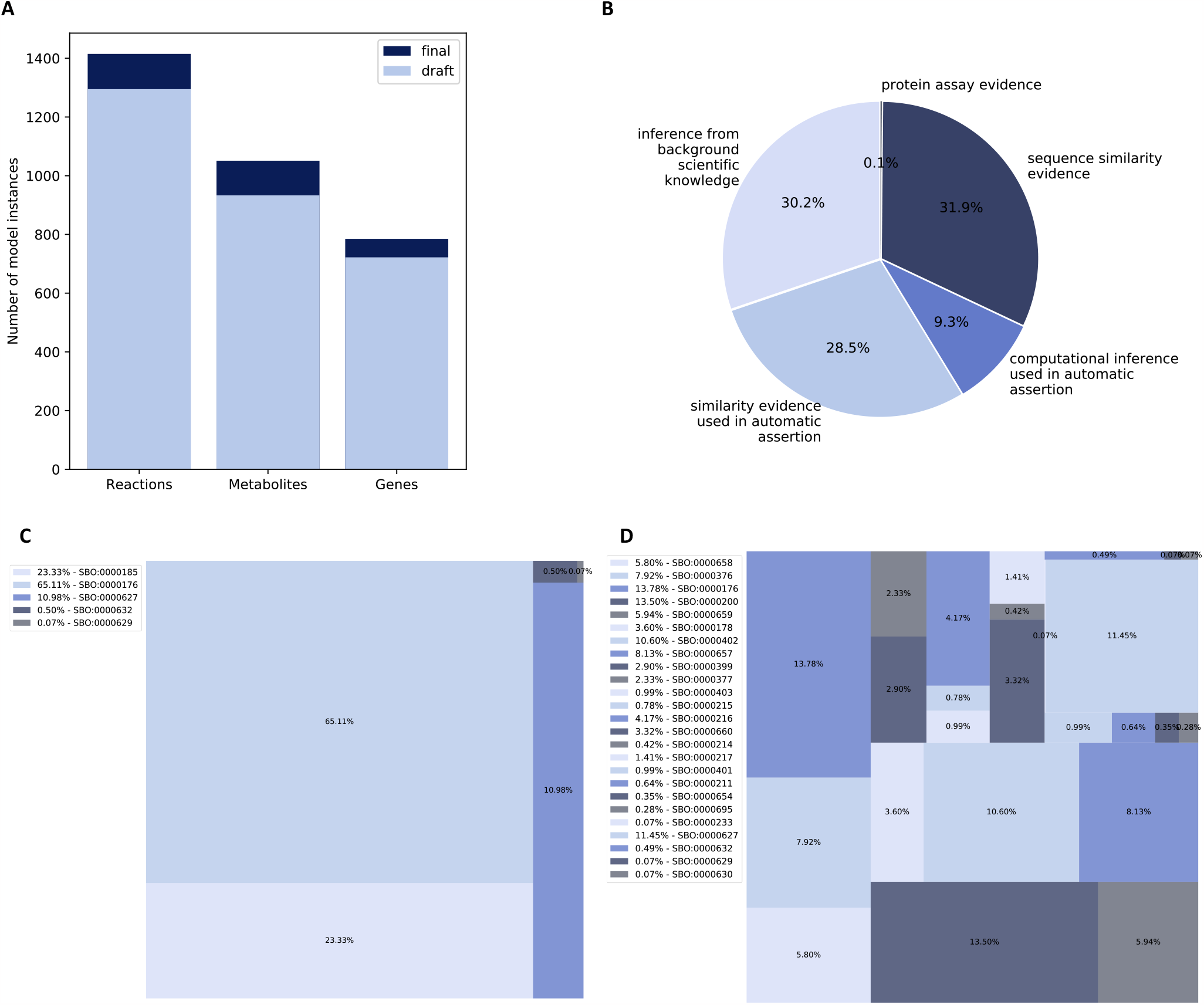
Properties of the network reconstructed for *S. epidermidis* ATCC 12228. **(A)** The initial draft network consisted of 1,295 reactions, 933 metabolites, and 722 genes. Further refinement and augmentation yielded the final metabolic model, comprising 1,415 reactions, 1,051 metabolites, and 785 genes. **(B)** To characterize the reactions, Evidence and Conclusion Ontology (ECO) terms were assigned based on the associated GPRs. The terms were allocated according to varying levels of evidentiary support. Notably, the term denoting inference from background scientific knowledge was assigned the lowest evidence level, while the term linked to protein assay evidence received the highest. **(C-D)** Coverage of Systems Biology Ontology (SBO) terms within the metabolic network before (C) and after (D) utilizing the SBOannotator^19^.

i. The cross-references to the nine databases are stored under the biological qualifier type `BQB_IS`.
ii. The ECO terms are stored under `BQB_IS_DESCRIED_BY`.
iii. Pathways associated with a reaction are saved with the biological qualifier type `BQB_OCCURS_IN`.

The metabolites and genes were annotated with twelve and three external databases, respectively, using the biological qualifier type `BQB_IS` (Table 1). The inclusion of ECO terms ensures a comprehensive understanding of evidence and assertion methodologies^36^, thereby facilitating robust quality control measures and evidence queries. The ECO term with the lowest evidence level is `ECO:0000001`, coding for inference from background scientific knowledge (Figure 2). This term was ascribed to 30.2 % of the biochemical reactions within the network. Notably, this percentage encompasses pseudo-reactions, such as exchanges, sinks, demands, and the biomass function. Within the group of 431 reactions associated with this ECO term, 170 pertained to pseudo reactions. The ECO term `ECO:0000251` denotes similarity evidence used in automatic assertion and was assigned to 28.5 % of all reactions. Moreover, the terms `ECO:0`(computational inference used in automatic assertion) and `ECO:0000044` (sequence similarity evidence) annotated 9.3 % and 31.9 % of all reactions, respectively. A minimal fraction (0.1 %) of reactions exhibits protein assay evidence, identified by the `ECO:0000039` term. Additionally, the SBOannotator was utilized to annotate the model with precise and descriptive SBO terms^19^ (Figure 2). Totally, 25 terms were incorporated describing classes of biochemical reactions and further model elements.

**Table 1.** Cross-references to various reaction, metabolite, and gene databases.

| Reactions | Metabolites | Genes |
| --- | --- | --- |
| BiGG Models<sup>20</sup> | BiGG Models<sup>20</sup> | NCBI protein<sup>21</sup> |
| MetaNetX<sup>22</sup> | MetaNetX<sup>22</sup> | UniProt<sup>23</sup> |
| KEGG<sup>24</sup> | KEGG<sup>24</sup> | KEGG<sup>24</sup> |
| ModelSEED<sup>25</sup> | ModelSEED<sup>25</sup> |  |
| BioCyc<sup>26</sup> | BioCyc<sup>26</sup> |  |
| UniPathway<sup>27</sup> | UniPathway<sup>27</sup> |  |
| Reactome<sup>28</sup> | Reactome<sup>28</sup> |  |
| IntEnz<sup>29</sup> | ChEBI<sup>30</sup> |  |
| Rhea<sup>31</sup> | HMDB<sup>32</sup> |  |
|  | InChIKey<sup>33</sup> |  |
|  | SABIO-RK<sup>34</sup> |  |
|  | LIPID MAPS<sup>35</sup> |  |

The final curated metabolic model was stored as a Systems Biology Markup Language (SBML) Level 3^37^ file. This format version supports the integration of various plugins, such as the `fbc` package^38^ and the `groups` package^39^, which are both enabled in *i*Sep23. The `groups` package facilitates the incorporation of additional information without impacting the mathematical interpretation of the model. We defined all pathways and subsystems identified from the Kyoto Encyclopedia of Genes and Genomes (KEGG) database^24^ as an individual group and added corresponding reactions as members. Overall, we added 99 distinct groups to the model that facilitate pathway-related analysis.

### Validation of the Metabolic Network

Besides the syntactic evaluation, data structure, and file format validation, the model also underwent assessment for its predictive value. A standard approach for such evaluations involves comparing simulation outcomes with empirical laboratory data. Given the adaptability of microbes to diverse environmental conditions, our focus was on investigating their growth behavior across various nutrient media. To enable comparability between simulation and laboratory results, ensuring that the simulated conditions represented those in the actual experiments was imperative.

#### Evaluation of Different Growth media

Therefore, we used chemically defined media for the growth simulation. In more detail, we utilized three synthetic minimal media: synthetic minimal medium (SMM)^40^, AAM^41^, and AAM-^42^. Developed initially to explore the metabolic requirements of *S. aureus*, these media definitions served as the basis for our simulations. We used the compound concentrations in the media definitions for the *in silico* simulations and tested whether our model exhibited positive growth with them. Furthermore, we extended our evaluations to include the widely used LB. Through a combination of *in silico* and *in vitro* experiments, we explored growth dynamics in the four media formulations both with and without d-glucose as the carbon source. This dual approach allowed us to measure growth in a simulated environment and in a real laboratory setting, providing a comprehensive validation of the model’s predictive performance under various nutritional conditions.

Figure 3 illustrates the growth behavior of *S. epidermidis* in various environments both *in silico* and *in vitro*. In minimal media where d-glucose serves as the sole carbon source, *S. epidermidis* could not exhibit growth. However, in the LB, *S. epidermidis* demonstrates the ability to utilize alternative carbon sources when glucose is absent. *In silico* simulations show growth in all tested minimal media, while *in vitro* experiments reveal no growth in AAM-, a medium lacking l-arginine. Comparative analysis of AAM-, AAM, and SMM highlights the absence of l-arginine in AAM-, a compound crucial for *S. epidermidis* growth. Prior studies have identified l-arginine auxotrophies in *Staphylococcus* species, including *S. epidermidis* ^44^. Despite reported l-arginine auxotrophy, the *S. epidermidis* strain ATCC 12228 harbors biosynthetic pathways for l-arginine via l-ornithine and l-glutamate, as reported in BioCyc^26^ and KEGG ^24^. In AAM-, l-glutamate is not provided as an amino acid, but it can be synthesized from l-proline, an amino acid present in the medium. The biosynthetic pathway is illustrated in Figure 4. All available l-proline is taken up and subsequently metabolized to, amongst others, l-glutamate, l-ornithine, and l-arginine. Each reaction in this pathway is supported by genetic evidence through a gene-reaction rule, commonly known as GPR.

**Figure 3.**
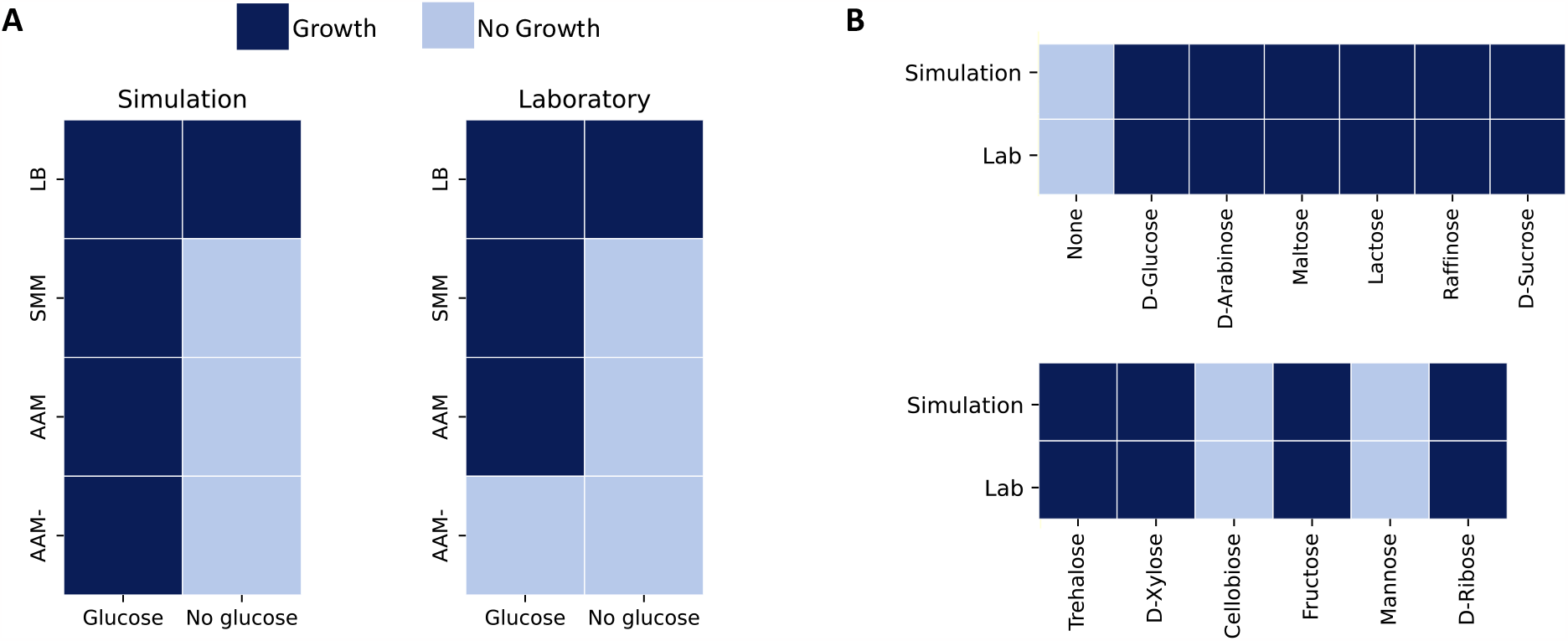
Growth phenotypes of *S. epidermidis* in different nutritional environments. **(A)** Evaluation of *S. epidermidis*’ growth encompassed various environmental conditions, including testing on complete LB and three minimal media formulations: SMM, AAM, and AAM-, both with and without d-glucose as a carbon source. The computational model successfully simulated growth across all media when glucose was the carbon source. However, the model predicted growth exclusively in the lysogeny broth (LB) without glucose. Experimental verification supported these findings, except growth on AMM-with d-glucose as a carbon source, where no growth was observed. **(B)** Analysis of growth in different carbon sources utilized SMM as the primary medium, wherein equimolar amounts of other sugars systematically replaced glucose. Simulation results closely paralleled laboratory findings, ensuring consistency across computational predictions and experimental outcomes.

**Figure 4.**
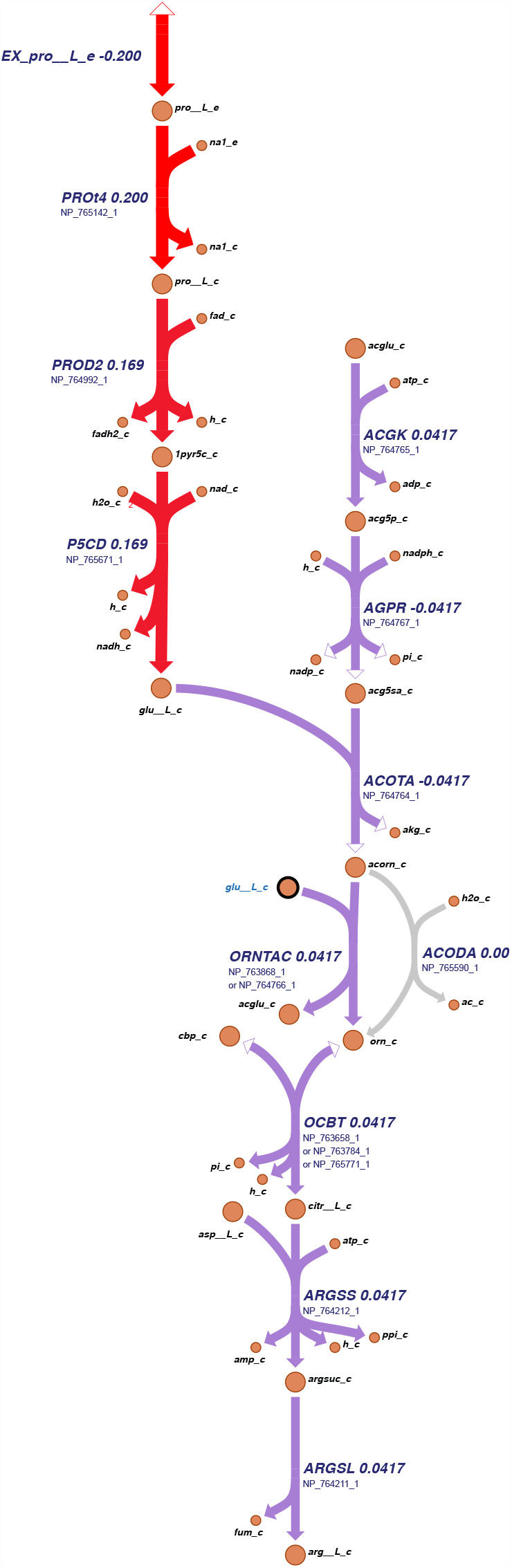
Biosynthetic pathway of l-arginine via l-glutamate and l-proline. All available l-proline is actively taken up and subsequently metabolized to various products, including l-glutamate, l-ornithine, and ultimately l-arginine. Genetic evidence supporting each reaction is provided in the form of a driving enzyme associated with a gene-reaction rule. The values assigned to these reactions correspond to the flux distribution in AMM-. The graphical representation of the metabolic map was generated using Escher^43^.

#### Growth in different carbon sources

In addition to evaluating *S. epidermidis*’s growth behavior in different media, we assessed the utilization of various carbon sources. This involved employing SMM and substituting d-glucose with alterative sugars in amounts adjusted for carbon content. A total of 12 different sugars were subjected to evaluation, as illustrated in Figure 3. Except for cellobiose and d-mannose, *S. epidermidis* demonstrated the capability to utilize all tested sugars as a carbon source, both through computational simulations (*in silico*) and laboratory experiments (*in vitro*). This consistency between model predictions and experimental observations lends robust support to the accuracy of the computational model.

## Discussion

Here, we present a manually curated GEM of *S. epidermidis* ATCC 12228, *i*Sep23. Literature-based corrections and meticulous manual curation ensured accurate representation of enzymatic reaction directions, essential for precise constraints during simulations. Overall, our model aligns with experimental data and offers a comprehensive platform for exploring *S. epidermidis*’s metabolic capabilities and behavior under diverse conditions. The inconsitency between the *in silco* and *in vitro* results reagrding the AAM-in the presence of glucose could be attributed to factors beyond the metabolic scope. For instance, non-metabolic factors could be regulatory mechanisms and Post-translational modifications. The observed discrepancy suggests a need for a more detailed understanding of the regulatory and metabolic factors influencing *S. epidermidis* growth in AAM-. Further experimental validation and exploration of regulatory mechanisms are crucial for resolving the observed differences between *in silico* predictions and experimental outcomes.

All in all, the refined network serves as a powerful tool for exploring *S. epidermidis*’s metabolic capabilities and behavior under diverse conditions. Future perspectives involve leveraging the model for targeted studies, such as investigating metabolic pathways, assessing the impact of genetic modifications, and exploring potential drug targets. The model’s compatibility with the `fbc` and `groups` packages in the SBML Level 3 Version 1^12^ format enhances its flexibility, enabling the integration of additional plugins for more intricate analyses. Including 99 distinct groups representing pathways and subsystems from the KEGG database provides a foundation for comprehensive pathway-related analyses. Altogether, *i*Sep23 aligns with experimental data and lays the groundwork for future investigations into the bacterium’s metabolism. Its accuracy, comprehensibility, and flexibility make it a valuable resource for advancing our understanding of microbial physiology and metabolic engineering applications.

## Materials and Methods

### Reconstructing the draft model of *S. epidermidis*

The reconstruction of the GEM is based on protocols described in previous studies^45, 46^. The fast and automated reconstruction tool CarveMe^47^ curates genome-scale metabolic models of microbial species and communities^47^. During the initial curation phase, a universal model was systematically compared to the annotated genome sequence of the species of interest, facilitating the construction of individual single-species metabolic models. In this study, we utilized CarveMe version 1.2.2 and the annotated genome sequence of *S. epidermidis* ATCC 12228 with the RefSeq^48^ accession ID `NC_004461.1` that covers the bacterial chromosome. Throughout the drafting process and subsequent model iterations, rigorous monitoring and benchmarking were conducted using MEMOTE ^15^. MEMOTE performs standardized metabolic tests across four key domains: annotation, basic tests, biomass reaction, and stoichiometry. The results are stored in a comprehensive report that includes the model’s overall performance assessed by a metric called MEMOTE score (denoted as a percentage with 100 %). A higher MEMOTE score correlates with enhanced annotation quality, greater consistency, and formal correctness of the model in SBML^49^ format. To refine the initial model automatically, the ModelPolisher^50^ was employed in a preliminary step. Leveraging the Biochemical, Genetical, and Genomical (BiGG) Models database^20^ identifiers of the model instances, the ModelPolisher systematically accessed the BiGG Models database, assimilating all available information for these instances into the network as annotations.

### Manual refinement of the draft metabolic network

In the initial draft model, a total of 63 reactions were identified as exhibiting mass and charge imbalances. To rectify these imbalances, an investigation into the connectivity of metabolites was conducted, focusing on identifying those frequently participating in reactions characterized by imbalances. To ensure the accuracy of the corrected model, the databases MetaNetX^22^ and BioCyc^26^ were browsed. These databases provided essential information about the correct charges and chemical formulas of the metabolites involved in the identified reactions, facilitating the precise adjustment of mass and charge imbalances within the model.

Additionally, the network constraints were carefully reviewed. Enzymes frequently act as catalysts in metabolic reactions. However, some enzymes effectively catalyze the reaction only in one direction. Consequently, it becomes imperative to impose constraints on the directionality of a given reaction. Instances where irreversible reactions are erroneously modeled as reversible, can result in an artificial expansion of the solution space within simulations. Conversely, misrepresenting reversible reactions as irreversible can unduly constrict the solution space, thereby precluding potential solutions. During our analysis, we systematically assessed various reaction directionalities and rectified any inaccuracies as necessary.

#### Detecting energy-generating cycles

GEMs with EGCs may harbor thermodynamically inaccurate cycles capable of generating energy without concurrent nutrient consumption^51^. These undesirable loops necessitate detection and subsequent elimination from the model. Fritzemeier et al. developed a systematic workflow for different energy metabolites. For each energy metabolite, a dissipation reaction was introduced into the model. After the imposition of constraints whereby all uptake rates were set to zero, an optimization process was conducted on the dissipation reaction. The presence of a non-zero flux following optimization serves as an indicator of the existence of EGCs within the model.

#### Including gene annotations

The software ModelPolisher^50^ was used to annotate the model instances. It is noteworthy, however, that this tool does not facilitate the annotation of model genes due to their strain-specific nature. We annotated the network genes using the associated National Centre for Biotechnology Information (NCBI) protein identifiers^21^. Notably, these gene identifiers underwent modifications during the reconstruction process due to the prokaryotic RefSeq genome re-annotation project^48^. To address this, we retrieved the updated NCBI protein identifiers from the NCBI database^21^. Subsequently, leveraging these novel protein identifiers in conjunction with the organism’s GenBank file^52^, we extracted the corresponding KEGG gene identifiers, which align with the organism’s locus tag and UniProt identifiers^23^. The integration of cross-references was executed as annotations using libSBML^53^. This comprehensive process ensures the accuracy and coherence of gene annotations within the model, thereby contributing to the reliability and accuracy of subsequent analyses.

#### Adding subsystems and groups

The reaction-associated pathways were retrieved using the annotated KEGG identifiers and the KEGG Representational State Transfer (REST) Application Programming transfer Interface (API). Subsequently, these pathways were incorporated as annotations utilizing the biological qualifier `BQB_OCCURS_IN`. Furthermore, the `groups` package was activated for enhanced functionality. Each identified pathway was integrated as a group, and the corresponding reactions as members.

#### Adding ECO and SBO terms

To enhance the model’s reusability, we incorporated ECO terms that annotate all metabolic reactions^36^. This ontology comprises terms and classes of the various evidence and assertion methods. These terms elucidate, for instance, the nature of evidence associated with a gene product or reaction, thereby facilitating robust model quality control. The assignment of a suitable ECO term to each reaction involved the extraction of GPRs. In instances where a reaction lacked a GPR, the term `ECO:0000001` was ascribed, denoting its inference from background scientific knowledge. Conversely, for all reactions with a GPR, the protein’s existence was reviewed in the UniProt database^23^. We distinguished the presence of proteins based on distinct categories, namely: (i) inferred from homology (`ECO:0000044`), (ii) predicted (`ECO:0000363`), (iii) evidence at the transcript level (`ECO:00000009`), or, or (iv) protein assay evidence. Genes not found in UniProt were assigned the term `ECO:0000251`, indicating the similarity evidence used in an automatic assertion. The relevant ECO term was incorporated as an annotation in instances where a biochemical reaction was associated with a GPR described by a single gene. In cases where the GPR involved multiple genes, the gene associated with the lowest evidence score was appended. All ECO terms were supplemented with the biological qualifier `BQB_IS_DESCRIBED_BY`.

The SBOannotator^19^ was employed to assign SBO terms to all reactions, metabolites, and genes within the metabolic network. These terms offer clear and unambiguous semantic information, delineating the type or role of each individual model component.

#### Elimination of redundant information

CarveMe stores the annotation information on model instances and cross-references to external databases within the notes field. However, the annotation field in the form of the controlled vocabulary (CV) terms is more appropriate for this information. Hence, we transferred all cross-references to the annotation field using the ModelPolisher^50^. Subsequently, to optimize file size and eliminate redundancy in information storage, the annotation information was systematically removed from the notes field.

#### Model extension

Model extension involved the integration of supplementary reactions sourced from established literature. The knowledge bases utilized for this purpose included BioCyc^26^, KEGG ^24^, and ModelSEED^25^. To identify relevant genetic information, locus tags from gene annotations were extracted and compared against the KEGG database. Reactions catalyzed by hypothetical enzymes were excluded from analysis. Candidate reactions were systematically cross-referenced with the BiGG ^20^ and ModelSEED databases, and were subsequently integrated into the network with BiGG identifiers and corresponding GPRs. If no entry in the BiGG database was specified, reaction identifiers from the source database were used.

### Evaluation and validation of growth capabilities

#### Different growth media

The growth behavior of *S. epidermidis* was assessed in three distinct synthetic minimal media initially formulated for investigating the metabolic requirements of *S. aureus*. These are the: (i) SMM^40^, (ii) AAM^41^, and (iii) AAM-^42^; a modified version of the AAM medium. The concentrations of the various components served as lower bounds for the corresponding exchange reactions of metabolites, as detailed in Table 2. In addition to the already provided salts and ions, we added minimal traces of zinc (`EX_zn2_e`), cobalt (`EX_cobalt2_e`), and copper (`EX_cu2_e`) to the simulated medium to enable growth. The lower bound of these reactions was set to −0.0001 mmol/(g_DW_ · h). Oxygen availability was defined by setting the lower bound of the exchange reaction to −20 mmol/(g_DW_ · h). The initial formulation of the three media involved the use of nicotinic acid. However, as nicotinic acid was substituted with nicotinamide in laboratory experiments, our simulated media also incorporated nicotinamide. In addition to the three minimal media, we tested *S. epidermidis*’s growth on the LB^47^. The lower bounds of the compounds’ exchange reactions listed in the LB were set to −10 mmol/(g_DW_ · h). All *in silico* simulations were evaluated with and without d-glucose as a carbon source.

**Table 2.** Testing growth of *S. epidermidis* in different synthetic minimal media. The values were set as lower bounds for the respective reactions and carry the unit mmol/(g_DW_ · h).

| Compound | Reaction ID | AAM | AAM- | SMM |
| --- | --- | --- | --- | --- |
| Sodium | EX_na1_e | -9.500 | -9.500 | -9.500 |
| Chloride | EX_cl_e | -12.522 | -12.522 | -12.522 |
| Potassium | EX_k_e | -3.140 | -3.140 | -3.140 |
| Magnesium | EX_mg2_e | -1.300 | -1.300 | -1.300 |
| Sulfate | EX_so4_e | -5.316 | -5.316 | -5.316 |
| Water | EX_h2o_e | -10.000 | -10.000 | -10.000 |
| Ammonium | EX_nh4_e | -4.000 | -4.000 | -4.000 |
| Calcium | EX_ca2_e | -0.022 | -0.022 | -0.022 |
| Phosphate | EX_pi_e | -0.140 | -0.140 | -0.140 |
| Iron | EX_fe2_e | -0.006 | -0.006 | -0.006 |
| Manganese | EX_mn2_e | -0.010 | -0.010 | -0.010 |
| Citrate | EX_cit_e | -0.006 | -0.006 | 0 |
| D-glucose | EX_glc__D_e | -5.000 | -5.000 | -5.000 |
| L-arginine | EX_arg__L_e | -0.125 | 0 | -0.125 |
| L-cysteine | EX_cys__L_e | -0.080 | -0.080 | -0.080 |
| L-leucine | EX_leu__L_e | -0.150 | -0.150 | 0 |
| L-glutamate | EX_glu__L_e | -0.250 | 0 | 0 |
| L-proline | EX_pro__L_e | -0.200 | -0.200 | -0.200 |
| L-threonine | EX_thr__L_e | -0.150 | 0 | 0 |
| L-valine | EX_val__L_e | -0.150 | 0 | 0 |
| L-phenylalanine | EX_phe__L_e | -0.150 | 0 | 0 |
| Nicotinamide | EX_ncam_e | -0.002 | -0.002 | -0.002 |
| Thiamin | EX_thm_e | -0.002 | -0.002 | -0.002 |
| Pantothenate | EX_pnto__R_e | -0.002 | -0.002 | -0.002 |
| Biotin | EX_btn_e | -0.0001 | 0 | 0 |
| Proton | EX_h_e | -0.140 | -0.140 | -0.140 |

#### Different carbon sources

Twelve different sugars were tested for their potential role as a carbon source: d-glucose, d-arabinose, maltose, lactose, raffinose, d-sucrose, trehalose, d-xylose, d-cellobiose, fructose, mannose, and d-ribose. For the growth simulations in different carbon sources, we used the SMM with nicotinamide instead of nicotinic acid as a basis (see Table 3). The concentrations reported in the medium were established as lower bounds for the simulation. The concentrations of the listed carbon sources were calculated to be equivalent in carbon content to the initial 5 g/L of glucose used in the defined SMM.

**Table 3.**
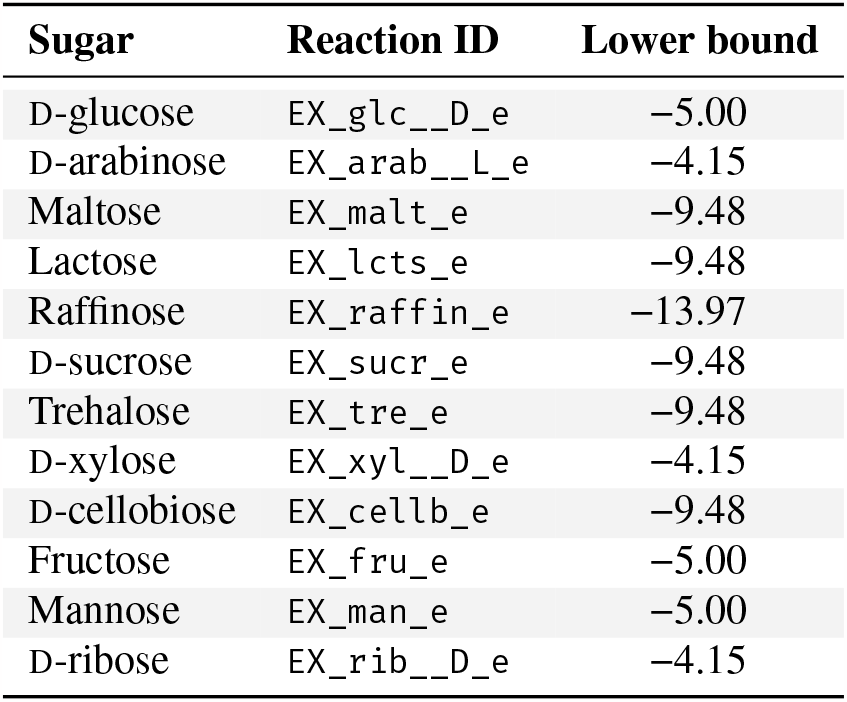
Testing of different carbon sources. Twelve different sugars were tested for their potential to serve as a carbon source in *S. epidermidis*. All values are given in mmol/(g_DW_ · h).

| Sugar | Reaction ID | Lower bound |
| --- | --- | --- |
| D-glucose | EX_glc__D_e | −5.00 |
| D-arabinose | EX_arab__L_e | −4.15 |
| Maltose | EX_malt_e | −9.48 |
| Lactose | EX_lcts_e | −9.48 |
| Raffinose | EX_raffin_e | −13.97 |
| D-sucrose | EX_sucr_e | −9.48 |
| Trehalose | EX_tre_e | −9.48 |
| D-xylose | EX_xyl__D_e | −4.15 |
| D-cellobiose | EX_cellb_e | −9.48 |
| Fructose | EX_fru_e | −5.00 |
| Mannose | EX_man_e | −5.00 |
| D-ribose | EX_rib__D_e | −4.15 |

### Laboratory validation

#### Media preparation

The minimal media AAM, AAM-, and SMM were prepared as carbon-source free base media following the methods provided by Machado et al. after omitting glucose as the default carbon source^40^. The carbohydrates to replace glucose as alternative carbon sources were dissolved in their respective base medium, and the resulting media were sterile filtered. Carbohydrates were obtained from Carl Roth (d-arabinose, d-glucose, trehalose, lactose, sucrose, raffinose), EMD-Millipore (fructose), Fluka (maltose, d-cellobiose), and Sigma Aldrich (mannose, d-ribose, d-xylose) in purity grades of *≥* 98 %. LB was prepared following the standard formulation of 10 g/L tryptone (MP Biomedicals), 10 g/L sodium chloride (Carl Roth), 5 g/L yeast extract (Carl Roth), and 5 g/L glucose when required.

#### Growth experiments

Cultures of *S. epidermidis* ATCC 12228 were initiated by inoculating overnight precultures in LB at 37 °C. Subsequently, primary cultures in LB were established from them and allowed to grow to an optical density (OD) at 600 nm (OD_600 nm_) of 0.5. Cell harvesting was achieved through centrifugation and two washes with the carbon-source-free medium. The cells were then resuspended to an OD_600 nm_ of 0.05 in media containing the respective carbon source. Growth was assessed by determination of the OD after a 24 h-incubation at 37 °C. Growth experiments were performed in at least three biological replicates in a 96-well plate format. OD measurements were performed with a Tecan Spark microplate reader.

## Data availability

Supplementary data are available along with this article. Additionally, *i*Sep23 is available at the BioModels Database^9^ as an SBML Level 3 Version 1^12^ file.

## Acknowledgments

This work was funded by the *Deutsche Forschungsgemeinschaft* (DFG, German Research Foundation) under Germany’s Excellence Strategy – EXC 2124 – 390838134 and supported by the Cluster of Excellence ‘Controlling Microbes to Fight Infections’ (CMFI). F.G. and A.D. is supported by the German Center for Infection Research (DZIF, doi: 10.13039/100009139) within the *Deutsche Zentren der Gesundheitsforschung* (BMBF-DZG, German Centers for Health Research of the Federal Ministry of Education and Research (BMBF)), grants № 8020708703 and № 8016708710. The authors acknowledge the support by the Open Access Publishing Fund of the University of Tübingen (https://uni-tuebingen.de/en/216529).

## Author contributions

**Conceptualization:** F.G and A.D

**Data curation:** N.L. and A.R.

**Formal analysis:** N.L., A.R., B.W., and A.G.

**Funding acquisition:** F.G. and A.D.

**Investigation:** N.L., A.R., and B.W.

**Methodology:** N.L., A.R., A.G., and B.W.

**Project administration:** F.G. and A.D.

**Resources:** F.G. and A.D.

**Software:** N.L., A.R., and A.G.

**Supervision:** F.G. and A.D.

**Validation:** F.G. and A.D.

**Visualization:** N.L. and A.R.

**Writing–original draft:** N.L., A.R. and B.W.

**Writing–review & editing:** N.L., A.R., F.G., and A.D.

## List of Abbreviations

API: Application Programming transfer Interface
BiGG: Biochemical, Genetical, and Genomical
BMBF: Federal Ministry of Education and Research (*Bundesministerium für Bildung und Forschung*)
BMBF-DZG: *Deutsche Zentren der Gesundheitsforschung*
CMFI: Controlling Microbes to Fight Infections
CV: controlled vocabulary
DFG: *Deutsche Forschungsgemeinschaft*
DZIF: German Center for Infection Research
ECO: Evidence and Conclusion Ontology
EGC: energy generating cycle
EMBL: European Molecular Biology Laboratory
FAIR: Findable, Accessible, Interoperable, and Reusable
GEM: genome-scale metabolic model
GPR: gene-protein-reaction association
KEGG: Kyoto Encyclopedia of Genes and Genomes
LB: lysogeny broth
MEMOTE: Metabolic Model Testing
NCBI: National Centre for Biotechnology Information
OD: optical density
REST: Representational State Transfer
SBML: Systems Biology Markup Language
SBO: Systems Biology Ontology
SMM: synthetic minimal medium

